# Targeting HIF-2α in Colorectal Cancer Reveals a Cholesterol Biosynthesis–Dependent Ferroptotic Vulnerability

**DOI:** 10.1101/2025.09.08.674939

**Authors:** Prarthana J. Dalal, Rashi Singhal, Boyan Hu, Drew Stark, Nicholas J. Rossiter, Chesta Jain, Peter Sajjakulnukit, Costas A. Lyssiotis, Yatrik M. Shah

**Author notes:** <u>Corresponding Author</u> Yatrik M. Shah, PhD 7712B Med Sci II 1137 E. Catherine St. Ann Arbor MI 48109-5622. Both authors contributed equally (Prarthana J. Dalal, Rashi Singhal).

## Abstract

Colorectal carcinoma (CRC) remains a major cause of cancer-related mortality, with rising incidence in individuals under 55, highlighting the need for novel therapeutic strategies. Hypoxia-inducible factor 2 alpha (HIF-2α) has been genetically validated as a critical driver of colorectal tumorigenesis, with intestinal epithelium–specific deletion in mice markedly reducing tumor formation. PT2385, a selective small-molecule HIF-2α inhibitor applied in renal cell carcinoma treatment, has not been evaluated in CRC. Here, we demonstrate that HIF-2α inhibition with PT2385 alone fails to suppress CRC growth in vitro under normoxic or hypoxic conditions and in xenograft models in vivo. To identify vulnerabilities induced by HIF-2α blockade, we performed an unbiased CRISPR metabolic screen. This revealed cholesterol biosynthesis as a critical dependency. Targeting this pathway with clinically approved statins (atorvastatin, pitavastatin, simvastatin) synergized with PT2385 to suppress CRC cell growth, reduce colony formation, and enhance cell death. Mechanistic studies show that combined HIF-2α and HMG-CoA reductase inhibition with statins promotes ferroptosis, characterized by increased lipid peroxidation and depletion of antioxidant metabolites. These effects are fully reversed by the ferroptosis inhibitor liproxstatin-1. Genetic knockdown of HIF-2α or HMG-CoA reductase recapitulated enhanced sensitivity to combination therapy. In vivo, co-administration of PT2385 and atorvastatin significantly reduced tumor growth and increased ferroptotic cell death in xenografts, confirming the mechanistic link. Collectively, these findings uncover a metabolic vulnerability of CRC to dual HIF-2α and cholesterol biosynthesis inhibition, supporting a clinically actionable strategy that leverages safe, FDA-approved statins to potentiate HIF-2α-targeted therapy.

**Significance:** Combined HIF-2α inhibition and cholesterol biosynthesis inhibition using statins reveals a metabolic vulnerability in colorectal cancer that enhances ferroptosis, thus offering a clinically actionable strategy for therapeutic intervention.

## Introduction

Colorectal carcinoma (CRC) is a common malignancy with an alarming increase in incidence and mortality among individuals below the age of 55. Therefore, there is an urgent need to develop new therapeutic strategies for advanced CRC.

CRC is characterized by significant tissue hypoxia arising from the dynamics of countercurrent exchange in blood flow, variations in oxygen demands, altered tumor metabolism, and proximity to an anoxic lumen. Hypoxia inducible factors (HIFs) transcriptionally orchestrate the cellular response to hypoxia and are involved in cell proliferation, survival, cell death, migration, and metabolism.

Specifically, HIF-2α plays an important role in CRC progression. HIF-2α expression is elevated in CRC compared with normal colonic epithelium and correlates with reduced survival [1].

Murine intestinal epithelium–specific HIF-2α deletion markedly decreases tumor formation [2]. Mechanistically, HIF-2α promotes colorectal carcinogenesis through recruitment of intratumoral neutrophils [3], dysregulation of iron homeostasis [4], potentiation of Yes-associated protein 1 activity [5], and activation of the cyclooxygenase 2 / prostaglandin E2 axis [6].

HIF-2α has recently emerged as a therapeutic target. PT2385 is a first-in-class, oral HIF-2α small molecule inhibitor. It was the first HIF-2α antagonist tested in humans and led to the development of belzutifan, an FDA approved compound [7]. PT2385 blocks HIF-2α dimerization with ARNT to prevent HIF-2α target gene transcription. Clinical development initially focused on renal cell carcinoma (RCC), where loss of the von Hippel-Lindau (VHL) tumor suppressor drives constitutive HIF-2α accumulation [8]. Given preclinical evidence for HIF-2α in murine CRC, we hypothesized that HIF-2α inhibition may represent a viable therapeutic strategy.

We show that HIF-2α inhibition alone does not suppress CRC in vitro or in vivo. We next performed an unbiased CRISPR metabolic screen and identified cholesterol biosynthesis as a critical vulnerability. Combination HIF-2α inhibition and 3-hydroxy-3-methylglutaryl-coenzyme A (HMG-CoA) reductase inhibition synergistically suppressed CRC growth and enhanced cell death. HMG-CoA reductase (HMGCR) is the rate-limiting enzyme in cholesterol biosynthesis and the pharmacologic target of statins, a safe and widely prescribed class of agents for cardiovascular disease [9].

Mechanistically, we demonstrate that the increased cell death observed with combination HIF-2α and HMGCR inhibition is mediated by ferroptosis. Ferroptosis is cell death characterized by intracellular iron increase, phospholipid oxidation, and loss of glutathione peroxidase 4 (GPX4) function [10]. While novel ferroptosis-inducing agents are under investigation as anti-cancer therapeutics [11], our findings highlight the potential of repurposing statins with HIF-2α inhibition for CRC therapy.

## Results

### HIF-2α inhibition does not affect CRC growth in vitro or in vivo

To assess the therapeutic potential of HIF-2α inhibition in CRC, we evaluated whether PT2385 altered CRC growth in vitro. Under normoxia (21% O₂), PT2385 had no effect on HCT116 or SW480 proliferation (Fig.1A). We next hypothesized that reduced oxygen availability might sensitize CRC cells to PT2385. However, even under hypoxia (2% O₂), PT2385 did not alter HCT116 or SW480 growth (Fig.1B).

**Figure 1.**
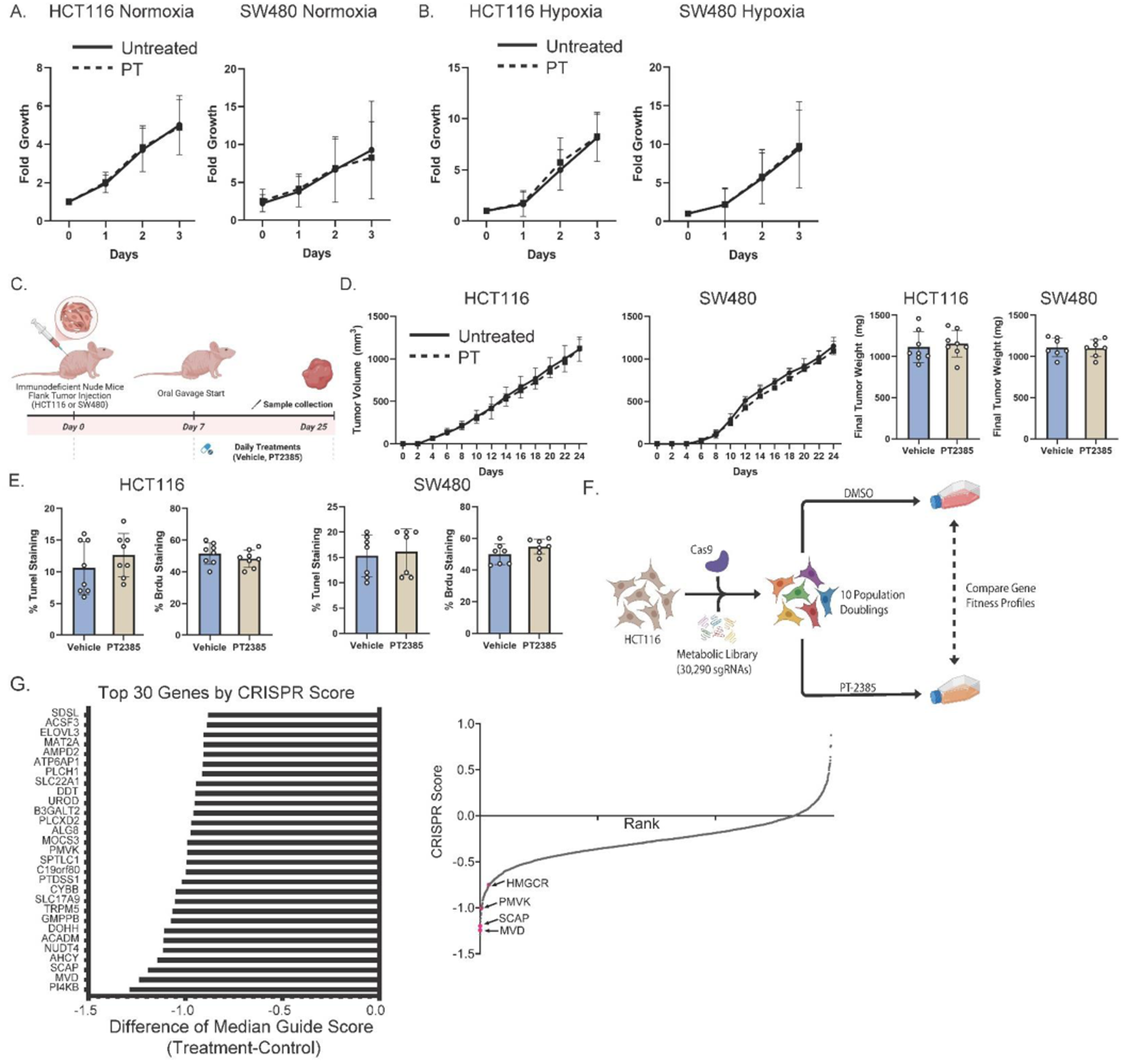
HIF2α inhibition does not alter CRC cell proliferation or tumor growth, but a CRISPR screen reveals specific metabolic dependencies. (**A**) Proliferation of HCT116 and SW480 colorectal carcinoma cells treated with PT2385 (PT) or vehicle under 21% O_2_. (**B**) Proliferation of HCT116 and SW480 cells treated with PT2385 or vehicle under 2% O_2_. (**C**) In vivo treatment schematic (**D**) Tumor growth kinetics and final tumor weights of HCT116 and SW480 xenografts in immunodeficient mice treated with either PT2385 or vehicle. (**E**) Quantification of cell death (TUNEL staining) and proliferation (BrdU staining) in harvested tumors. (**F**) Schematic of metabolic CRISPR screen in HCT116 cells comparing vehicle and PT2385 treated groups. (**G**) List of top 30 genes identified and figure of cholesterol biosynthesis pathway hits highlighted. Data is presented as mean ± SEM. Cell work was performed in triplicates and repeated three times. Mouse work was performed with 8 mice.

We next implanted HCT116 and SW480 cells into immunodeficient mice and administered PT2385 or vehicle (Fig.1C). PT2385 did not alter tumor growth kinetics or final tumor volume (Fig.1D). Cell death (TUNEL) and proliferation (Brdu) were unaffected in HCT116 and SW480 (Fig.1E).

To identify metabolic dependencies arising from HIF-2α inhibition, we performed an unbiased metabolic CRISPR screen (Fig.1F). Gene fitness profiles were compared between vehicle- and PT2385-treated cells. Key metabolic essentialities included phosphatidylinositol 4-kinase beta (PI4KB), mevalonate diphosphate decarboxylase (MVD), and adenosylhomocysteinase (AHCY) (Fig.1G). Cholesterol biosynthesis genes including sterol regulatory element binding protein cleavage activating protein (SCAP), phosphomevalonate kinase (PMVK), and HMGCR were also identified (Fig.1G).

### Combination HIF-2α inhibition and HMGCR inhibition suppresses CRC growth and enhances cell death

Selective small molecule inhibitors combined with PT2385 were tested to assess specific targets identified in our CRISPR screen. IN-10, a PI4KB inhibitor, did not impact HCT116 or SW480 proliferation with PT2385 (SFig.1A). 6-fluoromevalonate (6FM), an MVD inhibitor, also failed to enhance PT2385 efficacy (SFig.1B). Finally, 3-deazaneplanocin A hydrochloride (DZNep), a dual EZH2 and AHCY inhibitor, also did not affect CRC proliferation with PT2385 (SFig.1C).

However, the CRISPR screen identified cholesterol biosynthesis as an overall essential pathway of interest under HIF-2α inhibition. To target this with clinical relevance, we employed statins.

Simvastatin (Fig.2A), atorvastatin (SFig.2A), and pitavastatin (SFig.2A) showed a synergistic growth suppression with PT2385 in HCT116 and SW480 cells. The half-maximal inhibitory concentration (IC₅₀) of simvastatin was also reduced when combined with PT2385 (SFig.2B). Colony formation assays confirmed the long-term impact of combined inhibition (Fig.2B and SFig.2C).

**Figure 2.**
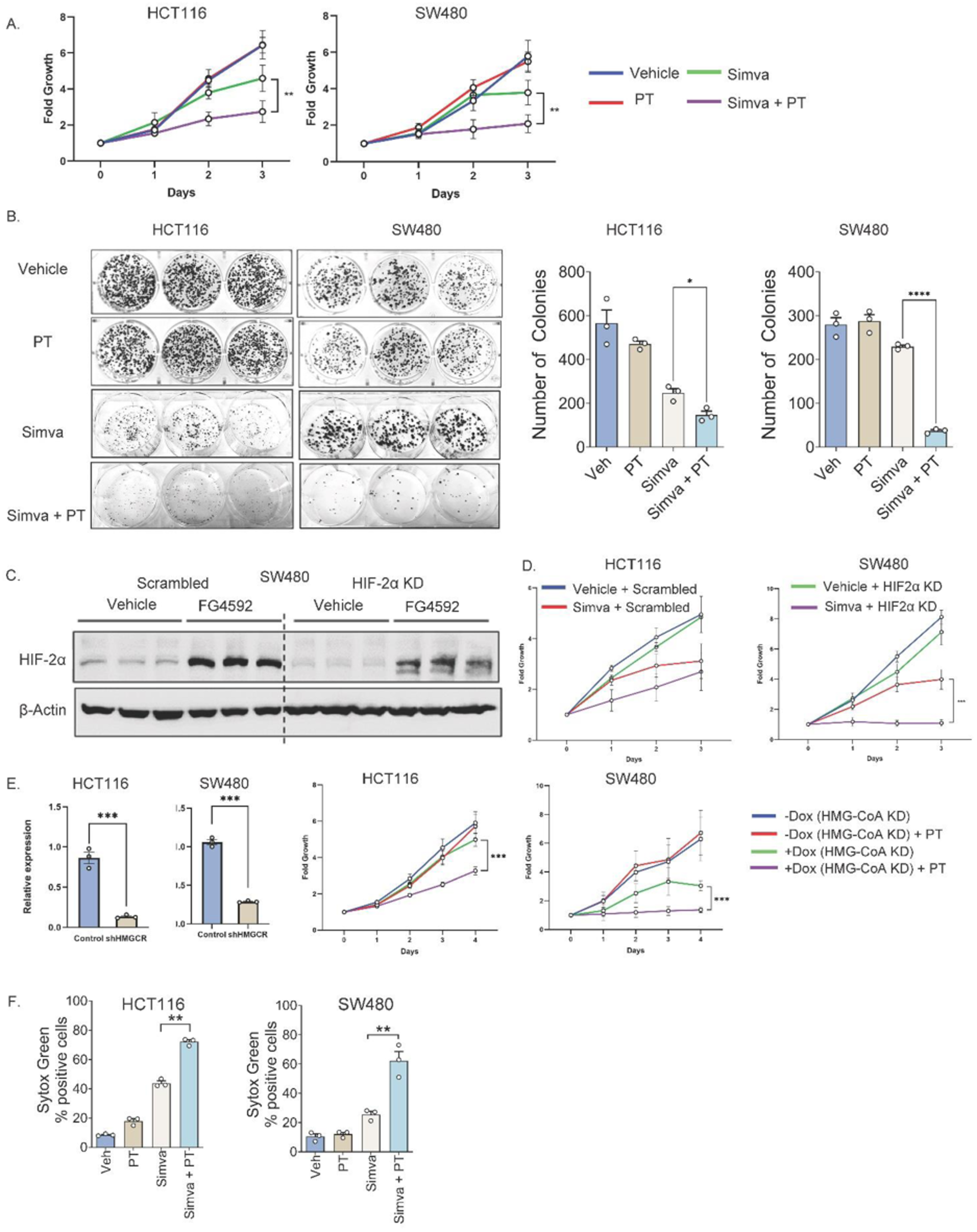
Pharmacologic and genetic inhibition of HIF-2α and HMGCR synergize to suppress CRC cell growth by inducing cell death. **(A)** Proliferation assays in HCT116 and SW480 cells treated with simvastatin (simva) alone or in combination with PT2385 (PT). **(B)** Colony formation assays demonstrate that simvastatin reduces colony number. This effect is further potentiated with PT2385. **(C)** SW480 cells transduced with shRNA targeting HIF2α show significant decrease in HIF2α protein level compared to vehicle in the presence and absence of FG4592, a hypoxia inducible factor prolyl hydroxylase inhibitor. **(D)** HCT116 and SW480 cells transduced with shRNA targeting HIF2α display decreased proliferation when treated with simvastatin. **(E)** HCT116 and SW480 cells transduced with shRNA targeting HMG-CoA reductase (HMGCR) show significant decrease in HMGCR expression by qPCR. Doxycycline-inducible, shRNA-mediated knockdown of HMGCR in HCT116 and SW480 cells decreases cell proliferation when combined with PT2385 (PT) treatment. **(F)** Sytox Green assays quantifying cell death in HCT116 and SW280 cells treated with simvastatin alone or in combination with PT2385 at 24 hours. Data are presented as mean ± SEM; *p<0.05, **p<0.01, ***p<0.001. Representative graphs shown. Pharmacologic proliferation assays performed in triplicates and repeated two times. Colony formation assays performed in triplicate. Genetic KD proliferation assays performed in triplicate. Cell death assays performed in triplicates and repeated two times.

We next validated these findings using genetic approaches. We previously generated and published HIF-2α knockdown (KD) in HCT116 cells [12]. To extend these observations, we established a second HIF-2α KD model in SW480 cells, which was confirmed by Western blot (Fig.2C). HIF-2α was knockout was success both in the presence and absence of FG4592, a hypoxia-inducible factor prolyl hydroxylase inhibitor which mimics the hypoxic response. These cells exhibited reduced proliferation when treated with simvastatin (Fig.2D), atorvastatin (SFig.2D), or pitavastatin (SFig.2D). Conversely, doxycycline-inducible HMGCR shRNA KD in HCT116 and SW480 was validated by qPCR (Fig.2E) and showed increased sensitivity to PT2385 (Fig.2E). Sytox Green assays also showed enhanced cell death, supporting that the synergistic growth suppression was driven by cell death rather than reduced proliferation (Fig.2F, SFig.2E).

### HIF-2α inhibition and HMGCR inhibition synergize through ferroptosis

To explore the mechanism driving cell death, we assessed a panel of cell death inhibitors. Inhibition of apoptosis or necroptosis did not decrease cell death following combination statin and PT2385 treatment (SFig.3A). Adding liproxstatin-1, a selective ferroptosis inhibitor, to combination statin and PT2385 treatment decreased the degree of cell death as assessed by Sytox Green assays (Fig.3A, SFig.4A). Lipid reactive oxygen species (ROS), a hallmark of ferroptosis, was also quantified. PT2385 plus statins significantly elevated lipid ROS and this effect was abrogated by liproxstatin-1 (Fig.3B, SFig.4B).

**Figure 3.**
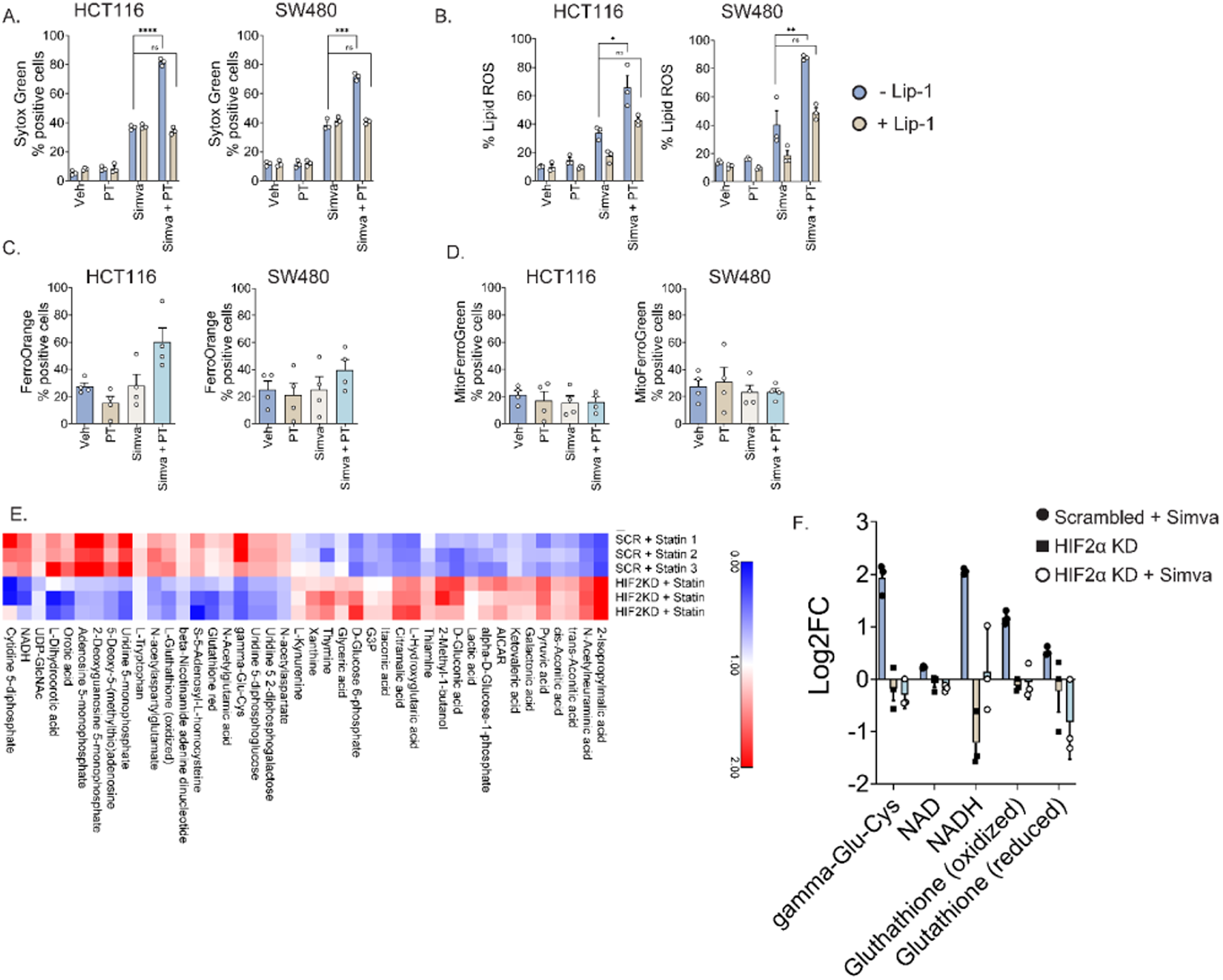
Combined statin and HIF2α inhibition triggers ferroptosis through increased lipid peroxidation and depletion of antioxidant metabolites in CRC cells. **(A**) Sytox Green assays in HCT116 and SW480 cells treated with simvastatin, PT2385, and liproxstatin-1 (selective ferroptosis inhibitor) demonstrate reduced cell death compared to simvastatin and PT2385 alone. **(B)** Combined simvastatin and PT2385 treatment significantly increases lipid ROS. This effect is rescued by the addition of ferroptosis inhibitor liproxstatin-1 (Lip-1) 1µM. **(C)** Cytosolic ferrous iron levels as measured by FerroOrange trended towards increase in HCT116 and SW480 cells with combined simvastatin (Simva) and PT2385 (PT) treatment compared to either agent alone. **(D)** Mitochondrial iron levels as measured by MitoFerroGreen remain unchanged. **(E)** LC/MS metabolomic analysis of HCT116 cells treated with simvastatin plus scrambled shRNA vs simvastatin plus HIF-2α shRNA reveals significant alterations in 42 metabolites. Combination treatment significantly impairs antioxidant capacity as demonstrated by γ-glutamylcysteine, NAD, NADH, reduced glutathione, and oxidative glutathione. **(F)** Metabolomic analysis of antioxidant capacity as demonstrated by γ-glutamylcysteine, NAD, NADH, reduced glutathione, and oxidative glutathione in HCT116 cells treated with HIF-2α shRNA vs HIF-2α shRNA and simvastatin. Data are presented as mean ± SEM; *p<0.05, **p<0.01, ***p<0.001. Cell death and lipid ROS assays performed in triplicates. Cytosolic and mitochondrial iron assays performed in triplicates and repeated four times. Metabolomics assay performed in triplicates.

**Figure 4.**
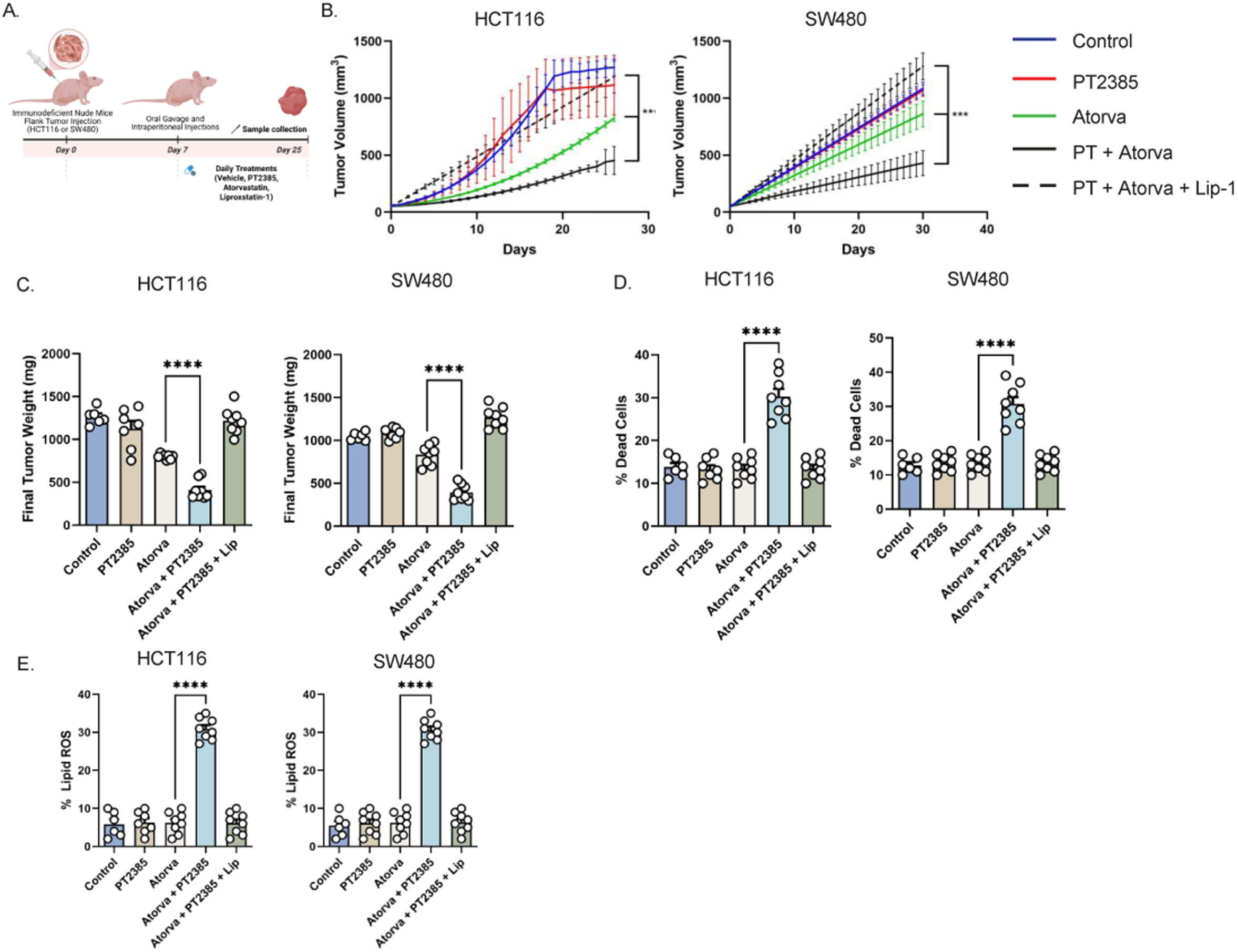
Dual HIF2α and HMG-CoA reductase inhibition with atorvastatin suppresses CRC tumor growth in vivo via ferroptosis-mediated cell death. **(A)** Experimental design schema **(B)** Tumor growth kinetics in HCT116 and SW480 xenografts treated with vehicle, PT2385, atorvastatin (Atorva), PT2385 (PT) plus atorvastatin, or PT2385 plus atorvastatin and the ferroptosis inhibitor liproxstatin-1. Combination PT2385 and atorvastatin treatment inhibited tumor growth, while liproxstatin-1 reversed this effect. **(C)** Final tumor weights show marked reduction with PT2385 plus atorvastatin, an effect reversed by liproxstatin-1 co-treatment. **(D)** Flow cytometry shows a significant increase in tumor cell death in the PT2385 plus atorvastatin group, an effect rescued by the addition of liproxstatin-1. **(E)** Lipid reactive oxygen species (ROS) were elevated in tumors treated with PT2385 plus atorvastatin but normalized with the addition of liproxstatin-1 further suggesting ferroptosis-specific cell death. Data are presented as mean ± SEM with 6-8 mice per group; *p<0.05, **p<0.01, ***p<0.001.

To further evaluate the role of iron in this process, we assessed intracellular ferrous iron using FerroOrange. Combined simvastatin and PT2385 treatment modestly increased cytosolic ferrous iron in HCT116 and SW480 (Fig.3C), whereas mitochondrial iron levels with Mito-FerroGreen were unchanged (Fig.3D).

To further investigate the metabolic basis of this, LC/MS metabolomic profiling of HCT116 cells treated with simvastatin and scrambled shRNA, HIF-2α KD shRNA alone, and simvastatin with HIF-2α KD shRNA was performed. Among 119 metabolites, 42 were significantly altered between simvastatin and scrambled shRNA and simvastatin and HIF-2α KD shRNA (p < 0.05; Fig.3E). Key oxidative stress intermediates including γ-glutamylcysteine (a glutathione precursor), NAD, NADH, and both oxidized and reduced glutathione were depleted in the combination group (Fig.3E). Oxidative stress intermediates were then compared between all three experimental groups (simvastatin and scrambled shRNA, HIF-2α KD alone, and HIF-2α KD with simvastatin treatment; Fig. 3F). The most pronounced differences are between the simvastatin and scrambled shRNA versus the simvastatin with HIF-2α KD groups. Overall, this suggests that HIF-2α inhibition diminishes antioxidant capacity to enhance susceptibility to lipid peroxidation and ferroptosis.

We also tested GPX4 protein levels in both HCT116 and SW480 CRC cells treated with vehicle control, PT2385, simvastatin, and combined PT2385 with simvastatin (SFig.4C). HIF-2α inhibition with PT2385 has no effect on GPX4 protein expression. Simvastatin alone decreases GPX4 protein expression. Combination treatment with PT2385 and simvastatin results in a more pronounced GPX4 protein decrease especially in SW480 cells. This suggests that while HIF-2α inhibition alone does not directly suppress GPX4, it sensitizes cells to mevalonate pathway inhibition. GPX4 is a selenoprotein and the mevalonate pathway is required for selenoprotein synthesis as it produces isopentenyl pyrophosphate (IPP), an intermediate essential for incorporating selenocysteine into selenoproteins. Ultimately these findings support that the mevalonate pathway is critical for selenoprotein synthesis in CRC and dual targeting with PT2385 and simvastatin can further compromise cellular GPX4-mediated defenses to enhance ferroptosis.

We also quantified lipid droplets in CRC cells treated with PT2385, statins (atorvastatin and simvastatin), and combination treatments (SFig. 5A). PT2385 does not appear to significantly induce lipid droplet biogenesis. Therefore, it is unlikely that PT2385 treatment alone predisposes to ferroptosis resistance due to sequestration of polyunsaturated fatty-acid (PUFA) rich lipids.

Finally, we also assessed whether PT2385 treatment decreases the output of the cholesterol biosynthesis pathway such that further inhibition with statins may be what drives the tumor growth defect. We performed quantitative PCR to assess the expression of various cholesterol biosynthesis genes and ultimately found that there is not statistically and biologically significant differences (SFig. 5B).

### HIF-2α inhibition and HMGCR inhibition decreases tumor growth through ferroptosis potentiation *in vivo*

For in vivo validation, we established xenograft models using HCT116 and SW480 cells and randomized mice into five treatment groups: vehicle, PT2385 alone, atorvastatin alone, PT2385 plus atorvastatin, and PT2385 plus atorvastatin with liproxstatin-1 (Fig.4A). Atorvastatin was chosen because it is a high intensity statin clinically preferred over simvastatin for its greater potency in lowering cholesterol [13]. Combination PT2385 and atorvastatin significantly reduced tumor growth kinetics (Fig.4B) and final tumor volume (Fig.4C). The addition of liproxstatin-1 fully reversed these effects, demonstrating ferroptosis dependence.

Flow cytometry revealed significantly increased proportions of dead cells in tumors treated with PT2385 plus atorvastatin, an effect abolished by liproxstatin-1 (Fig.4D). Similarly, lipid ROS levels were elevated in tumors receiving combination treatment, but this was also reversed by liproxstatin-1 (Fig.4E). This demonstrates that dual HIF-2α and HMGCR inhibition synergistically suppresses CRC growth in vivo through ferroptosis.

## Discussion

Our study demonstrates that unlike RCC, pharmacologic HIF-2α inhibition alone does not impair CRC growth. However, concurrent HIF-2α and cholesterol biosynthesis inhibition with statins produces a synergistic anti-tumor effect by inducing ferroptosis, highlighting tumor type–specific differences in HIF-2α dependency.

Given prior evidence of the contribution of HIF-2α to colorectal tumorigenesis, we initially expected PT2385 to suppress CRC growth [3–6]. However, PT2385 had no effect on CRC proliferation in vitro or in vivo. One explanation is that complete genetic loss of HIF-2α during tumor initiation fundamentally alters tumor biology, whereas acute pharmacologic inhibition in established tumors may be insufficient to disrupt compensatory pathways. Additionally, small-molecule antagonists such as PT2385 block HIF-2α/ARNT dimerization but may not fully recapitulate the broad transcriptional and metabolic consequences of HIF-2α deletion. These distinctions suggest that pharmacologic inhibition may unmask context-dependent vulnerabilities rather than phenocopying genetic ablation, leading us to investigate metabolic liabilities of HIF-2α inhibition.

An unbiased metabolic CRISPR screen identified cholesterol biosynthesis as a top vulnerability under HIF-2α inhibition. However, pharmacologic inhibition of a several top hits such as PI4KB, MVD, and AHCY did not further sensitize CRC cells to HIF-2α inhibition. This may be because genetic knockout via CRISPR may provide a more complete loss of function while pharmacologic inhibition may result in only partial suppression allowing residual protein activity to mask the sensitization phenotype. Additionally, variable drug bioavailability or specificity could dilute the inhibitor effect. Ultimately, we prioritized pharmacologic strategies directed at HMGCR due to their translational relevance.

Functional pharmacologic and genetic validation showed that HMGCR inhibition synergized with HIF-2α inhibition across multiple CRC cell lines. This effect was consistent across several statins, demonstrating a class effect rather than a drug-specific property. Dual inhibition promoted ferroptosis. Combination treatment increased cell death and lipid peroxidation, both effects reversed by the ferroptosis inhibitor liproxstatin-1. Metabolomic profiling demonstrated depletion of key antioxidant metabolites and impaired redox buffering, aligning with prior reports that statins decrease GPX4 expression and disrupt antioxidant pathways to sensitize cells to lipid peroxidation [14].

Interestingly, these findings appear paradoxical with our prior work, as well as independent studies from another lab, showing that HIF-2α inhibition increases ferroptosis resistance [4, 15]. This is further evidenced by inability to enhance cell death by combining a ferroptosis inducer, RSL3, with HIF-2α inhibition alone (data not shown). This may be explained by the metabolic rewiring that occurs with dual inhibition of both HIF-2α and cholesterol biosynthesis pathways specifically. HIF-2α inhibition alone decreases cellular antioxidants, lipid peroxidation capacity and confers ferroptosis resistance. However, simultaneous cholesterol biosynthesis inhibition appears to push cells past a metabolic threshold. Here antioxidant reduction by HIF-2α inhibition cooperates with statin-induced GPX4 impairment to sensitize cells to ferroptosis. This highlights the context-dependent effects of HIF-2α inhibition on ferroptosis. Furthermore, statins may also block the synthesis of coenzyme Q10 (CoQ10, ubiquinone) [16]. CoQ10 is essential to limit oxidative stress and lowering CoQ10 can further increase susceptibility to reactive oxygen species.

Beyond increasing sensitivity to oxidative stress and synergizing with GPX4 impairment, other possibilities for why cholesterol biosynthesis inhibition and HIF-2α inhibition synergize to induce CRC cell death include modulating lipid storage and endoplasmic reticulum (ER) stress. In RCC HIF-2α dependent lipid storage through perilipin 2 (PLIN2) allows tumor cells to buffer against sources of ER stress [17]. There may be a similar mechanism in CRC cells where inhibiting HIF-2α can alter ER lipid content, disrupt ER homeostasis, and trigger the unfolded protein response which in combination with statin induced alterations in oxidative stress trigger ferroptosis.

Clinically, belzutifan (MK-6482) is the next-generation HIF-2α inhibitor derived from PT2385 [7]. Although we employed PT2385 due to commercial availability, prior mouse studies have shown that PT2385 and belzutifan display equivalent efficacy and target engagement [18]. Thus, our findings are directly relevant to belzutifan. Nonetheless, future studies employing belzutifan with statins will be essential. Our data provides the first evidence that statins may sensitize CRC to HIF-2α inhibition, revealing a synthetic lethal interaction. It is also important to consider the effects of these drugs on non-cancer cells. Statins may lead to myalgias, while the most common side effect of HIF-2α inhibitors include anemia. Notably the most prominent side effects of these two drugs are largely non-overlapping, reducing the potential for compounded toxicity. Therefore, while both agents require appropriate clinical monitoring their combined use is likely tolerable for normal tissues. Given the widespread availability, safety, and low cost of statins, this approach could have significant translational potential.

## Materials and Methods

### Cell Lines and Reagents

HCT116 and SW480 cells (ATCC) were cultured in DMEM with L-glutamine, D-glucose, and sodium pyruvate, 10% heat-inactivated fetal bovine serum (FBS), and 1% antibiotic-antimycotic mix. IN-10 and DZNep were purchased from MedChemExpress. 6FM, Ferroorange (Cat# SCT210) MitoFerroGreen (Cat# SCT262) was purchased from MilliporeSigma. Atorvastatin, pitavastatin, simvastatin, Z-Vad, necrostatin-1, and FG4592 were purchased from Cayman Chemical (Cat# 10493, 15414, 10010344, 15294). Sytox Green was purchased from Thermo Fisher Scientific (Cat# S7020).

### Animals and Treatments

Animal procedures were approved by the University of Michigan Institutional Animal Care and Use Committee (PRO00011805). Six- to eight-week old male and female immunocompromised nude mice were inoculated with 2 million HCT116 or SW480 cells implanted as one tumor into the lower flank. Daily treatments began day 7 after visible tumors appeared. Oral gavage vehicle was PEG 400:Kolliphor (7:3). Intraperitoneal injection vehicle was sterile saline. PT2385 was given as oral gavage at 30mg/kg, atorvastatin as oral gavage at 10mg/kg, and liproxstatin as intraperitoneal injection at 10mg/kg.

### Western Blotting

Whole-cell lysates were prepared as previously described [19]. Primary antibodies including HIF-2α antibody (Novus Biological, Cat # NB100-122), GPX4 antibody (Protein Tech, Cat# 67763-1), and β-Actin antibody (Protein Tech, Cat # 66009-1) were used at 1:1000 dilution. Horseradish peroxidase-conjugated secondary antibodies were used at 1:2000 dilution and immunoblots were developed using Chemidoc imaging system.

### Growth Assays

Cell growth was measured with the Cytation 5 Imaging Multi-Mode reader with attached BioSpa (Agilent BioTek). Cells (∼500/well) were plated in 96-well plates. Baseline images were acquired after 24 hours and treatments were then immediately applied. Cell numbers were quantified every 24 hours and normalized to baseline values.

### CRISPR Screen and Analysis

The human CRISPR metabolic gene knockout library was a gift from David Sabatini (Addgene, 110066) [20]. Briefly ∼3,000 metabolic enzymes, small molecule transporters, and metabolism-related transcription factors are targeted. There are ∼30,000 sgRNAs corresponding to approximately 10 sgRNA/gene in addition to 500 control sgRNAs. 75 x 10^6^ HCT116 cells were seeded at 5 x 10^5^ cells/ml in six-well plates. Each contained 3mL DMEM, 10 µg/ml polybrene, and the CRISPR screen library virus. Target was to achieve 1,000 fold representation of each sgRNA in the library. Spinfection was performed by centrifugation at 1,200 *g* for 45 min at 37 °C. After 24 hours, the medium was replaced. Cells incubated for an additional 24 hours before puromycin selection (1 µg/mL). After three days of puromycin selection, cells were treated with DMSO or 10 µM PT-2385. Cells were passaged every 3-4 days. Following 14 doublings, genomic DNA was isolated from 15 million cells in each condition (DNeasy kit, Qiagen).

sgRNA inserts were PCR amplified using Takara Ex Taq polymerase, purified, and sequenced (NextSeq Illumina). Reads were mapped to the sgRNA reference library, allowing only exact matches, and the abundance of each sgRNA was quantified. For each sgRNA, a differential score was calculated as the log2 fold change in abundance between DMSO- and PT-2385 treated samples. Gene-level differential scores were determined by averaging the scores of all sgRNAs targeting the same gene. The gene score for each condition was defined as the median log2 fold change in abundance between the PT-2385 and vehicle treated populations.

### Generation of shRNA Cell Lines

Constitutively active genetically modified cell lines were generated using the commercially available human EPAS1 GIPZ lentiviral shRNA system targeting the open reading frame (Horizon). Doxycycline-inducible cell lines used the pLKO.1-Tet On system (Addgene, 21915) [21]. shRNA sequences were cloned into pLKO.1 backbones, sequence validated and made into lentivirus with the University of Michigan Vector Core. Cells were transfected through spinfection at 900 *g* for 1 hour in the presence of polybrene 10 µg/mL. After 24 hours the media was changed and cells allowed to recover for 24 hours before adding puromycin 2 µg/mL.

shRNA Primer Sequences:

HCT116 HMGCR FWD: CCGGCAACAGGTCGAAGATCAATTTCTCGAGAAATTGATCTTCGACCTGTTGTTTTT

HCT116 HMGCR REV: AATTAAAAACAACAGGTCGAAGATCAATTTCTCGAGAAATTGATCTTCGACCTGTTG

SW480 HMGCR FWD: CCGGTAGCCGTTAGTGGTAACTATTCTCGAGAATAGTTACCACTAACGGCTATTTTT

SW480 HMGCR REV: AATTAAAAATAGCCGTTAGTGGTAACTATTCTCGAGAATAGTTACCACTAACGGCTA

### Flow Cytometry for Sytox Green, FerroOrange, and Mito-FerroGreen

Cells were plated in six-well plates and treated for 16 hours. Mito-FerroGreen (5 µM) and FerroOrange (1µM) were added for 30 minutes at 37 °C. Cells were then washed and processed for flow cytometry. For Sytox Green assays cells were plated at 100,000 cells/well density in a 24 well plate. After seeding for 24 hours treatments were applied for an additional 24 hours. To harvest cells media was aspirated, cells washed once with PBS and then scraped off the plate. Cells were resuspended in PBS with 1µM working solution of Sytox Green dye. Cell pellet was incubated for 45 minutes at 37 degrees, washed with PBS and then processed for flow cytometry

### Lipid ROS Measurement

HCT116 or SW480 cells (1 x 10^6^) were seeded in 12-well plates. After 24 hours cells were treated as indicated for 16 hours. Cells were harvested, washed, and incubated with 5 µM C11-BODIPY (Thermo Fisher) for 30 minutes at 37 °C. A similar incubation period was used for *in vivo* studies and assessment of lipid ROS from murine tumors. Flow cytometry quantified fluorescence intensity. 20,000 cells minimum were analyzed per condition using FlowJo software.

### Lipid Droplet Staining and Microscopy

Ibidi µ-Slide 8 well high (Ibidi, 80807) were coated with 10 µg/cm^2^ human fibronectin (Sigma-Aldrich, FC010) following vendor recommendation. 15,000 SW480 and 10,000 HCT116 cells were then seeded per well and incubated at room temperature in the biosafety cabinet for 30 minutes before moving to the incubator. The next day, culture medium was replaced with medium containing 0.2% DMSO, 10 µM PT2385, 5 µM Atorvastatin, 3 µM Simvastatin, PT2385 and Atorvastatin combined, PT2385 and Simvastatin combined, or 200 µM Oleic Acid (Cayman Chemical, 24659) complexed with fatty acid free bovine serum albumin (Goldbio, A-421-10) at a 5:1 molar ratio in culture medium [22]. The following day, cells were washed once with staining medium (phenol red free DMEM with high glucose (Thermo Scientific, 31053028), supplemented with 4 mM L-glutamine (Research Products International, G36040) and 1 mM pyruvate (Agilent technologies 103578-100)). Cells were then incubated with 2 µg/mL Hoechst 33342 (Invitrogen H3570) and 1 µg/mL BODIPY 493/503 (Cayman Chemical, 25892) dissolved in staining medium for 25 minutes at 37 °C and 5% CO_2_. One untreated well per cell line was stained with Hoechst alone as an unstained control for the BODIPY dye. Cells were then washed once with staining medium without dyes and staining medium was replaced with staining medium supplemented with 10% FBS and 1% antibiotic-antimycotic mix before imaging.

Cells were imaged on a Zeiss LSM 980 Airyscan 2 confocal with spectral detection configured for multiplexing with a 405 nm laser line for imaging the Hoechst dye and a 488 nm laser line for imaging the BODIPY stain. The microscope was configured to maintain 37 °C and 5% CO_2_ with humidity and was stabilized for 30 minutes prior to imaging. Images were taken with a 63x oil objective with a 1.4 numerical aperture at 2x field of view. 35 z-stacks were taken for each image at the cell center point. Image post-processing was performed with Zen 3.4 software (Blue edition).

### Lipid Droplet Quantification

For HCT116 cells, 2 images were taken for the DMSO and unstained control conditions and at least 3 images were taken for all other conditions. Images were first configured as maximum intensity projections, and the maximum pixel value in the BODIPY channel for the unstained control images was subtracted from the BODIPY channel for all other images. Then, a manual threshold value was applied to both the BODIPY and the Hoechst channels across conditions to generate binary masks for each channel. Cells were manually segmented in each image by drawing ROIs around the nuclear and BODIPY masks for each cell. The area of the BODIPY mask was then calculated for each ROI.

For SW480 cells, 2 images were taken for the unstained control condition and at least 3 images for all other conditions. Images were analyzed the same as for HCT116 cells, except for thresholding was determined by FIJI using the automatic threshold calculation with the MaxEntropy algorithm. All image analysis was performed in FIJI [23].

BODIPY mask area/cell was analyzed and plotted using R v4.3.2 [24] with the dplyr [25] and ggplot2 [26] packages. A one way ANOVA with a post-hoc Tukey’s HSD test was performed to determine statistical significance.

### Colony Formation Assay

Cells were plated in biological triplicates in six-well plates at 500 cells per well. Media and treatments were replenished every 5 days. Assays concluded at 14 days by fixing cells with cold 10% buffered formalin for 10 minutes and staining with 1% crystal violet and 10% methanol solution for 30 minutes. Colonies were manually counted via a study blinded observer.

### Metabolomics

Cells were plated at 1 million/well in 6 well plates. After 24 hours, cells were treated for 24 hours. Metabolite isolation was performed using 80% methanol. Protein concentration for normalization was determined by processing a parallel well. Cell solutions were lyophilized using a SpeedVac concentrator. Metabolite pellets were resuspended in 50:50 methanol/water mixture for liquid chromatography (LC) – mass spectrometry (MS). Samples were analyzed on an Agilent 1290 Infinity II Bio LC ultra-high performance liquid chromatography system paired with an Agilent 6470 QQQ. Data were collected, processed, and analyzed using published parameters [27, 28]. Each metabolite abundance level was median-centered across all samples and log2-transformed to yield relative abundance for inter-group comparison. Statistical significance was determined by a one-way ANOVA with a significance threshold of p < 0.05.

### Statistics

Unpaired *t* tests determined statistical differences between 2 groups. A *p* value of less than 0.05 was considered statistically significant. All statistical tests were carried out using Prism software (GraphPad) or Excel.

**Supplemental Figure 1.**
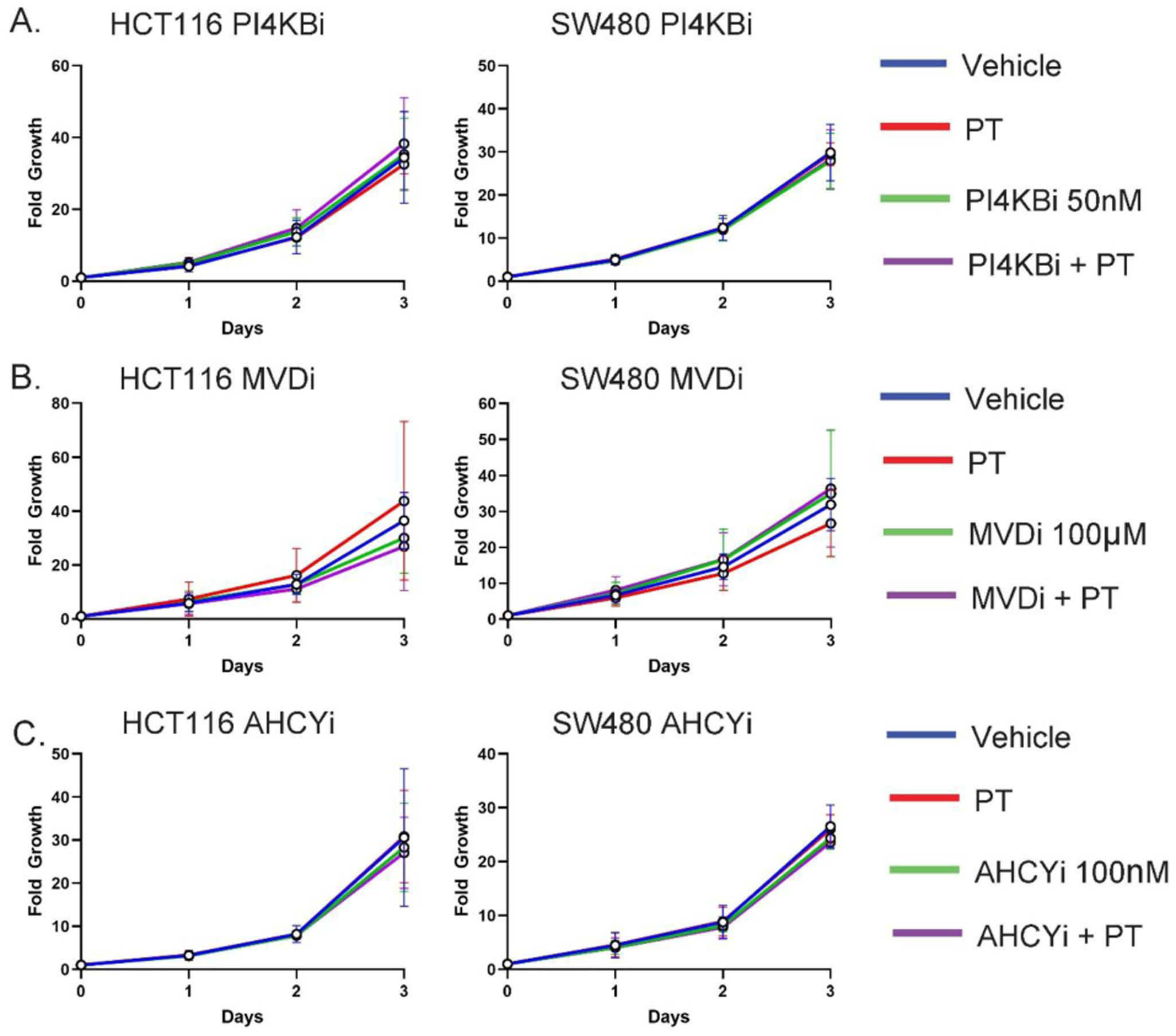
Metabolic vulnerabilities in PI4K, MVD, and AHCY do not sensitize CRC cells to HIF2α inhibition. **(A)** Proliferation of HCT116 and SW480 cells treated with PI4KB inhibitor IN-10, either alone or in combination with PT2385 (PT). **(B)** Proliferation of HCT116 and SW480 cells treated with MVD inhibitor 6-fluoromevalonate (6FM), alone or in combination with PT2385. **(C)** Proliferation of HCT116 and SW480 cells treated with dual EZH2 and AHCY inhibitor, 3-deazaneplanocin A hydrochloride (DZNep). Data are presented as mean ± SEM. Representative graphs shown. Proliferation data performed in triplicates and repeated 1-3 times.

**Supplemental Figure 2.**
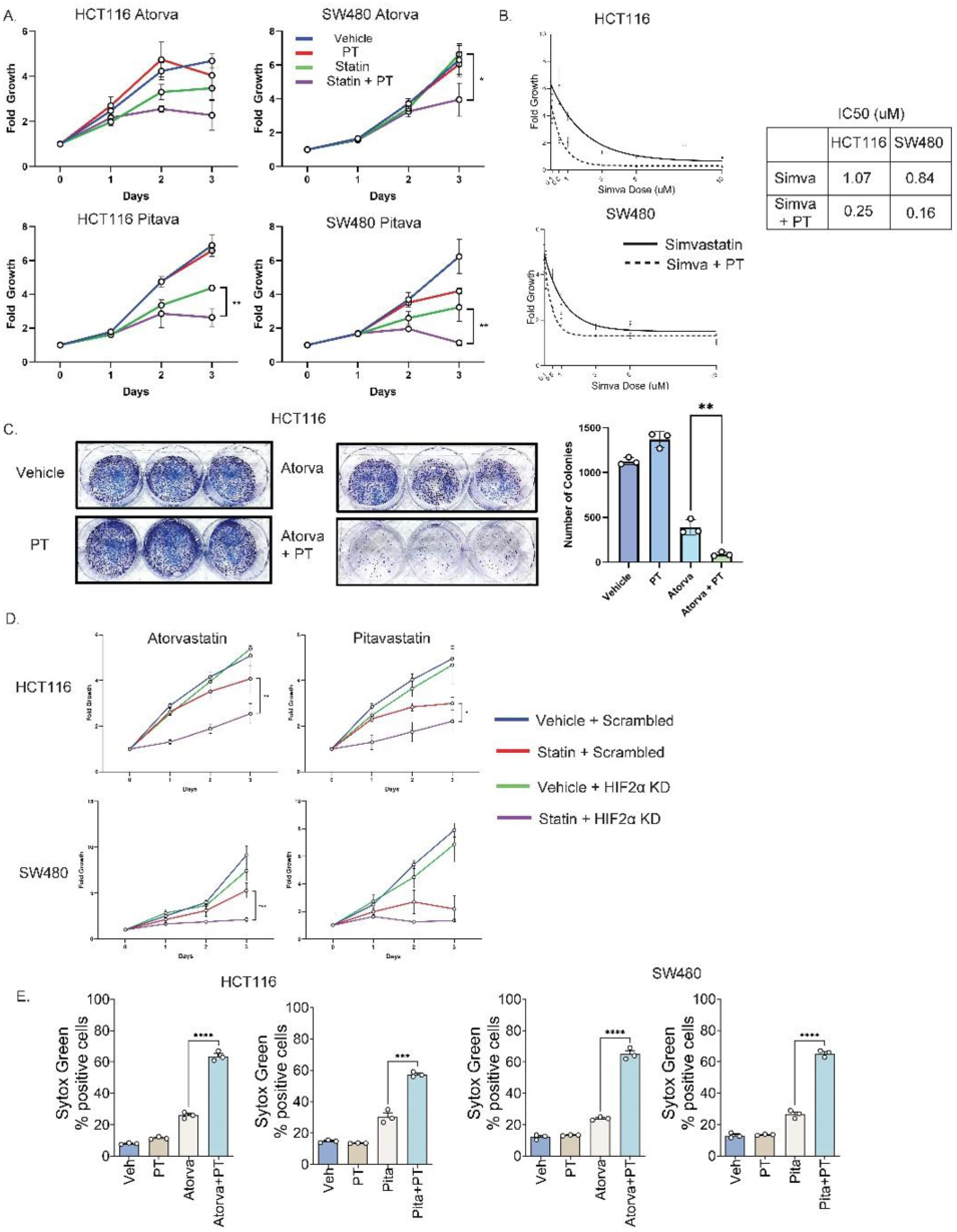
Atorvastatin and pitavastatin demonstrate similar effects in suppressing CRC and inducing cell death as simvastatin when combined with HIF-2α inhibition. **(A)** Proliferation assays in HCT116 and SW480 cells treated with atorvastatin (atorva) and pitavastatin (pitava) either alone or in combination with PT2385 (PT). **(B)** The half maximal inhibitor concentration (IC_50_) of simvastatin is reduced in the presence of PT2385. **(C)** Colony formation assays demonstrate that atorvastatin reduces colony number. This effect is further potentiated with PT2385. **(D)** HCT116 and SW480 cells transduced with shRNA targeting HIF2α display decreased proliferation when treated with atorvastatin or pitavastatin. **(E)** Sytox Green assays quantifying cell death in HCT116 and SW280 cells treated with atorvastatin or pitavastatin alone or in combination with PT2385 at 24 hours. Data are presented as mean ± SEM; *p<0.05, **p<0.01, ***p<0.001. Representative graphs shown. Proliferation data performed in triplicates and repeated 1-2 times. Half maximal inhibitor concentration performed in triplicates. Colony formation assay performed in triplicates. Cell death assays performed in triplicates.

**Supplemental Figure 3.**
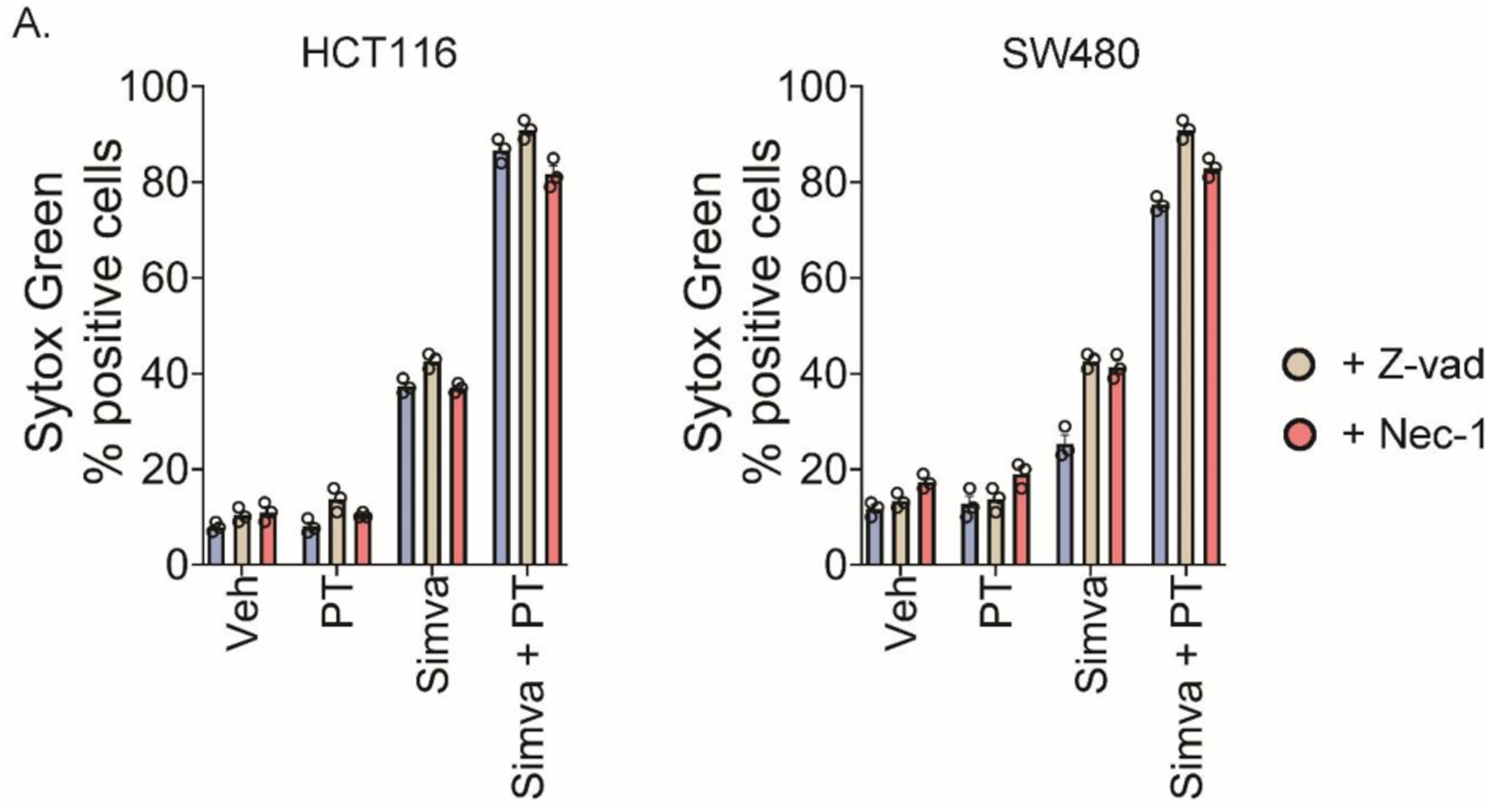
Inhibiting apoptosis and necroptosis cell death pathways do not reverse cell death mediated by combination simvastatin and HIF-2α inhibition. **(A**) Combined simvastatin (simva) and PT2385 (PT) treatment increases cell death as measured by Sytox Green. This effect is not reversible by the addition of a pan-caspase inhibitor blocking apoptosis-mediated cell death (Z-vad, 10 µM), or a RIP1 inhibitor blocking necroptosis-mediated cell death (Necrostatin-1, Nec-1, 10µM). Data are presented as mean ± SEM. Representative graphs shown. Cell death assays performed in triplicates and repeated two times.

**Supplemental Figure 4.**
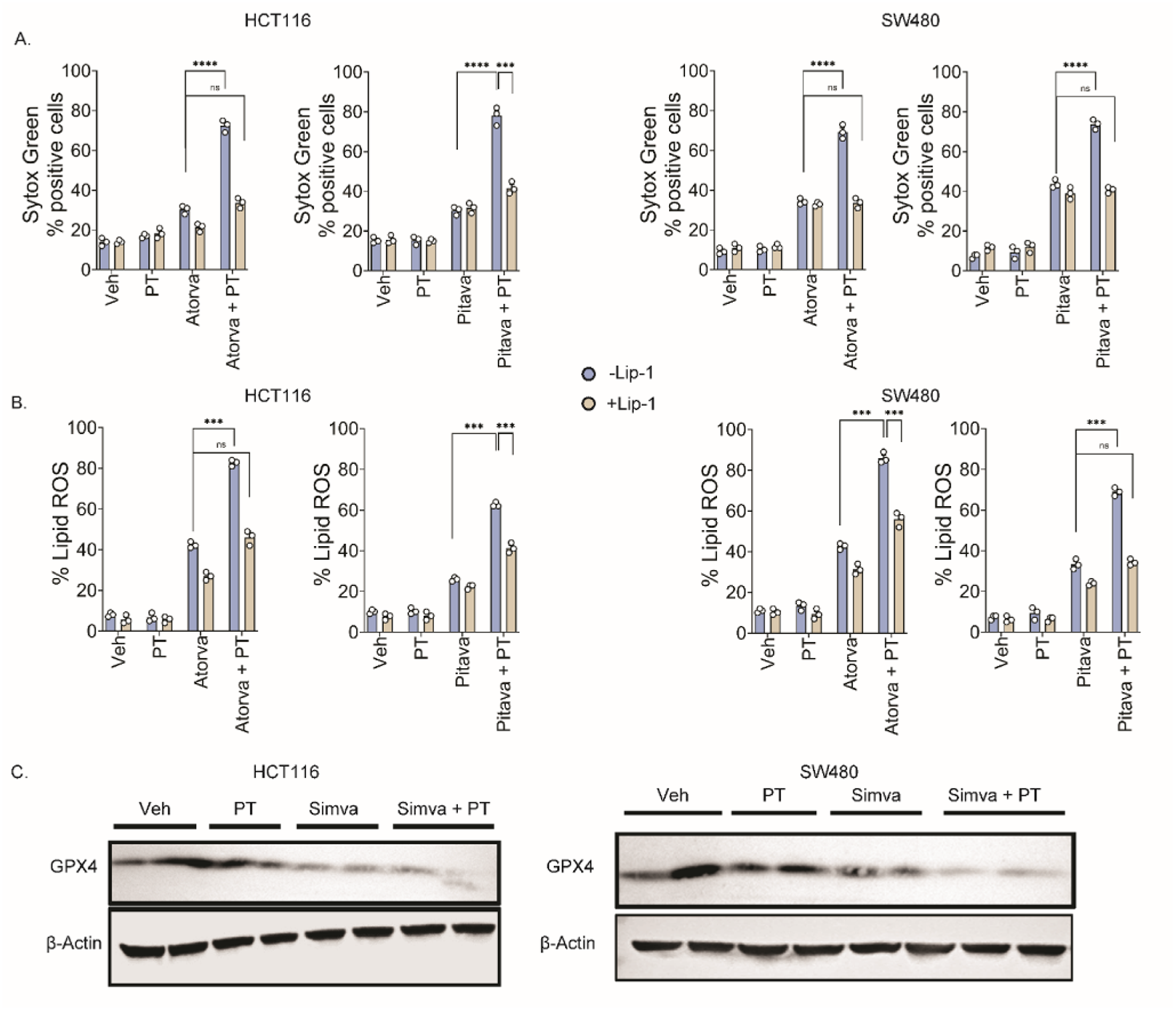
Increased cell death and lipid ROS generation caused by atorvastatin or pitavastatin in combination with HIF-2α inhibition is reversible with ferroptosis inhibitor liproxstatin-1. **(A**) Combined atorvastatin (atorva) or pitavastatin (pitava) and PT2385 (PT) treatment significantly increases cell death as measured by Sytox Green. This effect is reversible with the addition of ferroptosis inhibitor liproxstatin-1, 1µM (Lip-1). **(B)** Combined atorvastatin or pitavastatin with PT2385 treatment significantly increases lipid ROS. This effect is rescued by the addition of Lip-1. **(C)** Combined simvastatin (Simva) and PT2385 (PT) treatment decreases GPX4 protein levels especially in the SW480 CRC cell line. Data are presented as mean ± SEM; *p<0.05, **p<0.01, ***p<0.001. Cell death and lipid ROS data performed in triplicates.

**Supplemental Figure 5.**
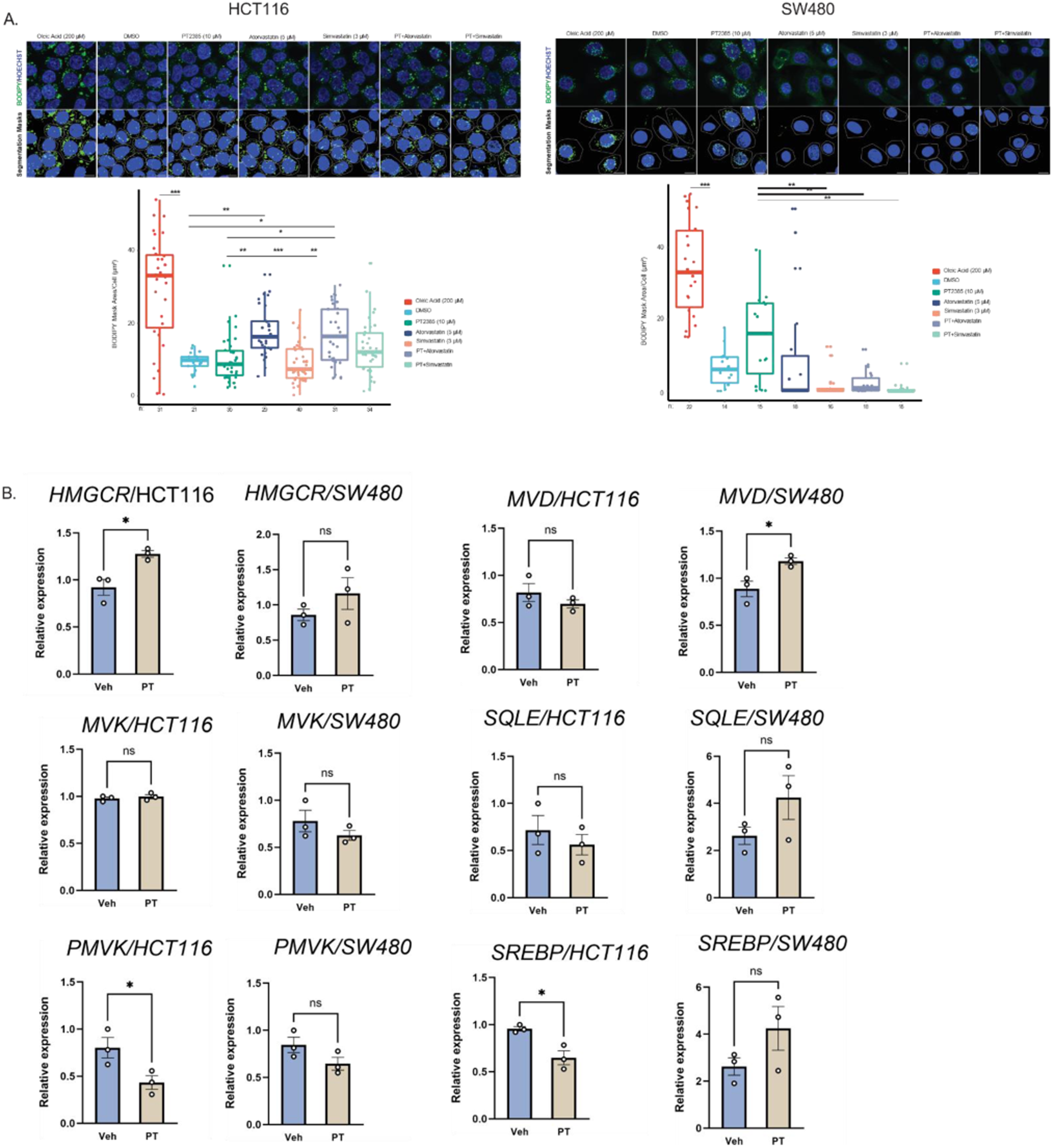
Lipid Droplet Quantification. **(A**) Lipid droplet area/cell was assessed in HCT116 and SW480 CRC cells using BODIPY 493/503. Cell segmentation and staining area was quantified across oleic acid, DMSO, PT2385, atorvastatin, simvastatin, and combination groups. Oleic acid significantly induces lipid droplet area. PT2385 treatment alone does not significantly alter lipid droplet area. * = p<0.05, ** = p<0.01, and *** = p<0.001. Scale bars = 10 µm **(C)** qRT-PCR Data on PT and cholesterol biosynthesis genes Data are presented as mean ± SEM; *p<0.05. qRT-PCR performed in triplicates.

